# The effects of cerebellar transcranial direct current stimulation on the cognitive stage of sequence learning

**DOI:** 10.1101/588012

**Authors:** Hannah K. Ballard, James R. M. Goen, Ted Maldonado, Jessica A. Bernard

## Abstract

Though the cerebellum has been previously implicated in explicit sequence learning, the exact role of this structure in the acquisition of motor skills is not completely clear. The cerebellum contributes to both motor and non-motor behavior. Thus, this structure may contribute not only to the motoric aspects of sequence learning, but may also play a role in the cognitive components of these learning paradigms. Therefore, we investigated the consequence of both disrupting and facilitating cerebellar function using high definition transcranial direct current stimulation (HD-tDCS) prior to the completion of an explicit motor sequence learning paradigm. Using a mixed within- and between-subjects design, we employed cathodal (n=21) and anodal (n=23) tDCS (relative to sham), targeting the lateral posterior cerebellum, to temporarily modulate function and investigate the resulting effects on the acquisition of a sequential pattern of finger movements. Results indicate that cathodal stimulation has a positive influence on learning while anodal stimulation has the opposite effect, relative to sham. Though the cerebellum is presumed to be primarily involved in motor function and movement coordination, our results support a cognitive contribution that may come into play during the initial stages of learning. Using tDCS targeting the right posterior cerebellum, which communicates with the prefrontal cortex via closed-loop circuits, we found polarity-specific effects of cathodal and anodal stimulation on sequence learning. Thus, our results substantiate the role of the cerebellum in the cognitive aspect of motor learning and provide important new insights into the polarity-specific effects of tDCS in this area.

**New & Noteworthy:** The cerebellum contributes to motor and cognitive processes. Investigating the cognitive contributions of the cerebellum in explicit sequence learning stands to provide new insights into this learning domain, and cerebellar function more generally. Using HD-tDCS, we demonstrated polarity-specific effects of stimulation on explicit sequence learning. We speculate that this is due to facilitation of working memory processes. This provides new evidence supporting a role for the cerebellum in the cognitive aspects of sequence learning.

## Introduction

The acquisition of motor skills is important for day-to-day functioning. Movement coordination is necessary for the operation of routine tasks, such as driving a car, riding a bike, playing a musical instrument, playing sports, etc. As such, the performance of these everyday tasks is dependent upon the ability to successfully learn and execute motor skills. Investigating the neural underpinnings of sequence learning, therefore, is essential for enhancing our knowledge and understanding of motor processes and allowing for effective protocols designed for improvement or rehabilitation of skilled motor behavior.

As suggested by Karni et al. (1998), sequence learning skills develop over time across distinct learning phases, beginning with fast, or early, learning and ending with slow, or late, learning. During the early learning phase, initial improvement in motor performance is observed. Additional improvement continues, and eventually, during the later phase of learning after extended practice, individuals reach a state of automaticity in performance. Distinct brain structures have been implicated in each sequence learning phase, contributing to the differences seen in skill progression (Doyon et al., 1997). The cerebellum is particularly active during the early learning phase when procedural memories are first being formed whereas the striatum takes over during later learning phases when performance improvements begin to level off (Ungerleider et al., 2002; Bernard et al., 2013; Doyon et al., 2018). Importantly, however, it may be that the cerebellum also contributes to the initial skill acquisition stages via its role in cognitive processing. A growing literature has indicated that the cerebellum contributes to cognition (e.g., Schmahmann and Sherman, 1998; Stoodley and Schmahmann, 2009; Stoodley, 2012), presumably through closed-loop circuits with the prefrontal cortex (cf. Strick et al., 2009).

In addition to the motor nodes associated with motor learning, non-motor regions of the brain also contribute to skill acquisition. As noted in a recent review by Doyon et al. (2018), cognitive processes play a role in the acquisition of new motor sequences. In functional neuroimaging investigations of motor adaptation, the dorsolateral prefrontal cortex was active in regions associated with cognitive function, specifically working memory, during the initial stage of learning (Anguera et al., 2010, 2012). Similarly, activation in the lateral prefrontal cortex, commonly implicated in cognitive processing, has also been demonstrated during explicit sequence learning (Honda et al., 1998; Sakai et al., 1998; Eliassen et al., 2001; Schendan et al., 2003; Aizenstein et al., 2004). Furthermore, working memory has been associated with the formation of motor chunks during the learning of a new sequence (Bo and Seidler, 2009). Given that the cerebellum plays a role in non-motor processing (e.g., Schmahmann and Sherman, 1998; Stoodley, 2012; Bernard and Seidler, 2013, 2014; Guell et al., 2018), in addition to contributing to the internal representation of motor responses during early sequence learning (Spampinato and Celnik, 2018), this structure may also be involved in the cognitive aspects of initial skill acquisition.

One way to investigate the role of the cerebellum in explicit motor sequence learning is to consider the effects of non-invasive brain stimulation on skill acquisition, using methods such as transcranial direct current stimulation (tDCS). With this technique, brain activity in a targeted area can be temporarily disrupted or facilitated, and the resulting impact on behavior can be observed (e.g., Nitsche and Paulus, 2000; Nitsche et al., 2008; Takano et al., 2011). At present, several studies have used tDCS to examine the role of different motor circuit nodes, including the cerebellum, in motor learning (e.g., Galea et al., 2011; Pope and Miall, 2012; Block and Celnik, 2013; Foerster et al., 2013; Shah et al., 2013; Hardwick and Celnik, 2014; Cantarero et al., 2015; Buch et al., 2017). These studies have established that the cerebellum plays a significant role in adaptation learning, but a clear understanding of the cerebellum’s involvement in explicit sequence learning, specifically, is still lacking. Therefore, though we know that the cerebellum is important for learning, investigating its contributions from a more cognitive circuit perspective with this particular type of task is necessary for extending current insights. Furthermore, high definition (HD) methods of tDCS (HD-tDCS) have now become available, allowing for better regional targeting of the stimulation. Notably, to our knowledge, HD-tDCS has not yet been employed to investigate the cerebellum in motor learning. This technique is particularly useful for investigating the cognitive contributions of the cerebellum in motor sequence learning as it allows us to better target the non-motor aspects of this structure.

Here, we targeted the lateral posterior cerebellum, primarily in the region of Crus I and Crus II, using cathodal and anodal HD-tDCS in two experiments. Notably, these regions are associated with prefrontal cortical networks in humans (Krienen and Buckner, 2009; O’Reilly et al., 2010; Salmi et al., 2010; Bernard et al., 2012, 2014) and in non-human primates (Kelly and Strick, 2003), in addition to being implicated across brain imaging studies of non-motor behavior (Stoodley and Schmahmann, 2009; E et al., 2014). Our goal was to investigate differences in motor sequence learning after purported inhibition with cathodal stimulation or excitation with anodal stimulation (Nitsche and Paulus, 2000), relative to sham. We predicted that cathodal stimulation targeting the lateral posterior cerebellum would negatively impact motor sequence learning relative to sham whereas anodal stimulation to the same cerebellar area would facilitate motor learning, due to the impact on cognitive function and associated cognitive circuits. These predictions are consistent with findings from recent sequence learning research using the traditional two electrode pad stimulation technique rather than the HD-tDCS approach employed here (Ehsani et al., 2016; Wessel et al., 2016; Shimizu et al., 2017). Thus, we conducted two independent experiments with a mixed within- and between-subjects design. For the within-subjects comparison, we employed cathodal and sham stimulation in Experiment 1 and anodal and sham in Experiment 2. We then compared across the cathodal and anodal groups in a between-subjects manner to get a complete idea of the effect of cerebellar stimulation on motor skill learning.

## Experiment 1

### Materials and Methods

#### Participants

Twenty-eight healthy young adults participated in the first experiment, which consisted of two HD-tDCS sessions (cathodal and sham) that were counterbalanced in order. Two of those individuals did not return for the second session of the study, one individual was not right-handed as per our screening criteria, and complete data was not collected from one individual due to technical difficulties. Additionally, three individuals were excluded from analyses for having task performance scores below three standard deviations from the group mean. This particular criterion was implemented as an *a priori* cut-off to exclude behavioral outliers. In investigating these outliers, the three individuals excluded had accuracy scores of 52% or lower. Indeed, one individual did not respond in time on as many as 88% of trials within a given block, suggesting that those excluded based on behavioral scores failed to comply with task instructions and may have performed poorly due to inattention or lack of motivation. As such, our final sample included twenty-one subjects (15 female, mean age 18.6 years ± 0.7 S.D., range 18-20 years). None of the subjects had any history of medical or neurological disease, and, as assessed by the Edinburgh Handedness Inventory, all were right-handed (mean score 96.8 ± 10.0 S.D.). Subjects were screened for exclusion criteria associated with tDCS (Nitsche et al., 2008), and none were taking medication that could potentially impact the central nervous system (e.g., psychiatric medications such as antidepressants, anxiolytics, neuroleptics, narcotics, stimulants, and analgesics). Subjects were recruited and scheduled through the Texas A&M University Psychology Subject Pool and received course credit for their participation. Participation was limited to individuals that were tDCS naïve. Prior to the initiation of study procedures, all subjects provided written informed consent. Our protocol was submitted to, and approved by, the Institutional Review Board at Texas A&M University.

#### Procedure

Participants underwent two sessions, exactly one week apart, in which they received both cathodal and sham tDCS in a within-subjects design. We used a single-blind procedure wherein participants were not told which condition they were receiving in a given session. The order of stimulation condition (cathodal or sham) was counterbalanced across all participants. Stimulation was administered with a Soterix MxN HD-tDCS system (Soterix Inc., New York, NY) using a montage of 8 stimulation electrodes. The electrode array and current intensities used for cathodal cerebellar stimulation are presented in Table 1 and the associated modeled current flow is depicted in Figure 1. Different intensities were used for each electrode location in order to effectively administer 2.0 mA of net current. The stimulation montage was obtained using HD-Targets software that provides specific electrode locations to more focally stimulate a targeted region of interest (Datta et al., 2016; Huang et al., 2017). In this case, our target was the right lateral posterior cerebellum. In determining the optimal stimulation montage, we chose a “spiral out” approach for this experiment, which is equivalent to cathodal stimulation in a traditional two electrode pad montage. A “spiral” approach is the terminology used to describe any montage containing multiple stimulation electrodes and distinguishes the direction of current flow. During each session, an initial stimulation of 0.1 mA was administered for one minute in order to allow the current to break through the scalp and establish an adequate connection. Once resistance levels of 50K or lower were achieved across all electrodes, the full stimulation period began, lasting 20 minutes. In the course of each condition, current gradually increased during the first 30 seconds. During active stimulation, the current remained steady for the entire span of the stimulation period. In the sham, or placebo, condition, subjects underwent the standard setup for HD-tDCS but only experienced current during the first and last thirty seconds of the stimulation period to allow for attenuation to occur and simulate the general experience associated with active stimulation. During this thirty-second period, the current slowly ramped up, and then quickly returned to zero. This sham procedure has been implemented in order to avoid the subjects’ detection of the stimulation condition as attenuation to the sensation of stimulation occurs relatively quickly under cathodal (or anodal) protocols.

**Table 1.** Electrode array and current intensities for cerebellar cathodal stimulation. *Location:* particular locations for electrode placement, determined by 10-20 system; *Current:* specific intensity of electrical current applied at a given location.

|  | <u>Location</u> | <u>Current</u> |
| --- | --- | --- |
| 1 | P2 | 0.1135 |
| 2 | PO10 | -0.1551 |
| 3 | O10 | -0.3618 |
| 4 | EX3 | 0.1956 |
| 5 | EX4 | -1.4832 |
| 6 | EX5 | 0.4066 |
| 7 | EX12 | 0.22 |
| 8 | EX14 | 0.7886 |
| C | NK1 | 0.2757 |

**Figure 1.**
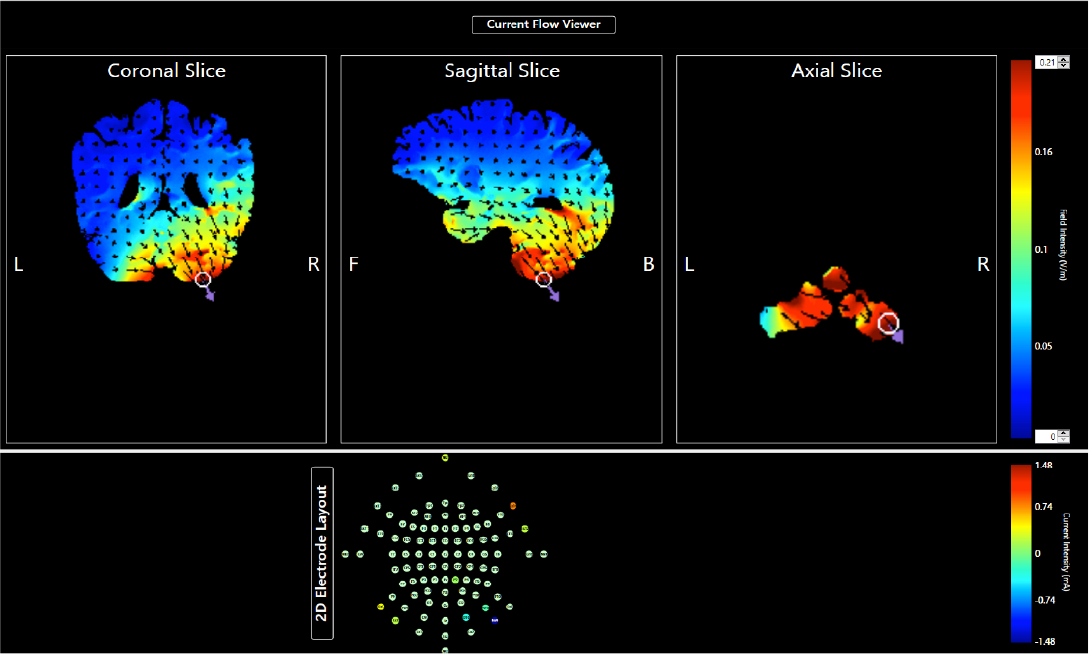
Modeled current flow for cerebellar stimulation using Soterix HD-Targets software. Color bar denotes field intensity (V/m) on a scale of 0 - 0.21 (bottom to top). *L*: Left; *R*: Right; *F*: Front; *B*: Back.

To assess the influence of tDCS on motor skill learning, we used an explicit motor sequence task modeled after Kwak et al. (2012). We chose to use this type of task in order to pinpoint the cognitive aspects of procedural learning, specifically working memory mediated strategies, that are primarily engaged in explicit paradigms. Indeed, investigations using functional magnetic resonance imaging (fMRI) have revealed greater activation in cognitive and working memory related areas during explicit, rather than implicit, learning (Jenkins et al., 1994; Grafton et al., 1995; Rauch et al., 1995; Schendan et al., 2003). In the task, participants were instructed to place their left middle, left index, right index, and right middle fingers over the numbers 1, 2, 3, and 4 on the computer keyboard, respectively. The task was bimanual in order to allow for translation to the MRI environment for follow-up work; this technique has been implemented in multiple other studies investigating sequence learning (Sun et al., 2007; Berner and Hoffmann, 2008; Bo and Seidler, 2009; Kwak et al., 2010, 2012; Bhakuni and Mutha, 2015). Participants were then presented with four rectangles in the center of the screen, corresponding to the instructed finger locations from left to right. In each trial, a different rectangle was shaded black, and participants were instructed to press the button matching the location of that shaded rectangle as quickly as possible. To account for baseline motor function, random blocks were included in between groups of sequence blocks throughout the task. Thus, each participant completed 6 random blocks with 18 trials each and 9 sequence blocks with 36 trials each for a total of 15 blocks and 432 trials (Kwak et al., 2012). Random blocks were delineated with an “R” prior to stimulus presentation, while sequence blocks were preceded by an “S”. Thus, the nature of the upcoming trials was announced before each block as the task was intended to be clearly explicit. Relatedly, our paradigm did not include extensive practice or subsequent retention sessions in order to avoid the recruitment of implicit learning processes in the development of automaticity. Each stimulus was presented for 200 milliseconds and participants were allowed 800 milliseconds to respond before the next stimulus appeared. The block design also mirrored that of Kwak et al. (2012) with the following order: R-S-S-S-R-R-S-S-S-R-R-S-S-S-R. To account for practice effects, participants were presented with a new sequence pattern during the second experimental session. Each sequence was comprised of 6 elements and did not contain any trills or sequential numbers. The following sequences were used in our design: 2-4-1-3-2-4 and 3-2-4-1-2-4; these sequences did not evidently differ in complexity as there was no significant difference in distribution between sessions, according to the Kolmogorov-Smirnov test (*D* = 0.182, *p* = 0.423).

#### Data Analysis

Performance on the motor sequence learning task was measured using average total accuracy (ACC) per sequence block and mean reaction time (RT) for correct responses during both random and sequence blocks. Accuracy is presented as the percent correct out of the total number of trials for each sequence block. Total trials included correct, incorrect (e.g., commission errors), and missed responses (e.g., omission errors). In order to remain consistent with the current sequence learning literature (Kwak et al., 2012) and factor in timing-specific effects on skill acquisition, we analyzed accuracy through a learning phase perspective. The first three sequence blocks during the task were concatenated and represented the early learning phase whereas the middle three blocks represented the intermediate phase, and the last three blocks represented the late learning phase, in the context of our study set-up. We chose to concatenate across sequence blocks in this manner to get at the more board changes in motor skill learning that may occur over the course of the task, as our paradigm was relatively brief. This approach was adopted instead of modeling across each individual block in order to simplify the analyses and avoid excessive follow-up comparisons that may complicate the interpretation of results, as other studies have similarly done (Bo and Seidler, 2009; Kwak et al., 2010, 2012). However, in the grand scheme of the sequence learning literature, our design still reflects an early learning paradigm that focuses on initial skill acquisition as extensive practice or subsequent experimental sessions using the same sequence were not included. In recognition of the overall brief nature of the task, we did not find it necessary to normalize scores relative to initial performance as baseline learning was not expected to influence later learning over the course of the task with only 9 sequence blocks, total, being executed. For RT, missed trials were excluded and we used correct trials only from both random and sequence blocks when calculating averages. Repeated-measures ANOVAs and paired sample *t-*tests were performed to investigate differences in performance on the motor sequence learning task between cathodal and sham stimulation. All analyses were conducted using SPSS Statistics.

### Results

#### Reaction Time

A 2 × 2 repeated measures ANOVA (block type by stimulation condition), collapsing across all random and sequence blocks respectively, revealed a significant main effect of block type, *F* (1, 20) = 108.23, *p* < .001, η^2^_p_ = .844, indicating that responses were significantly faster on sequence blocks compared to random, as expected. However, there was no significant block type by stimulation condition interaction, *F* (1, 20) = .144, *p* = .71, η^2^_p_ = .001, and no significant main effect of stimulation condition, *F* (1, 20) = 1.92, *p* = .18, η^2^_p_ = .087, indicating that RT did not differ between cathodal and sham stimulation. The difference in RT between block types indicates that learning took place as anticipating the upcoming key presses allowed for faster responses (Figure 2). In fact, *t*-tests reveal a largely significant difference in reaction time between the first (S2) and last (S14) sequence blocks (*p* < .001), further indicative of learning.

**Figure 2.**
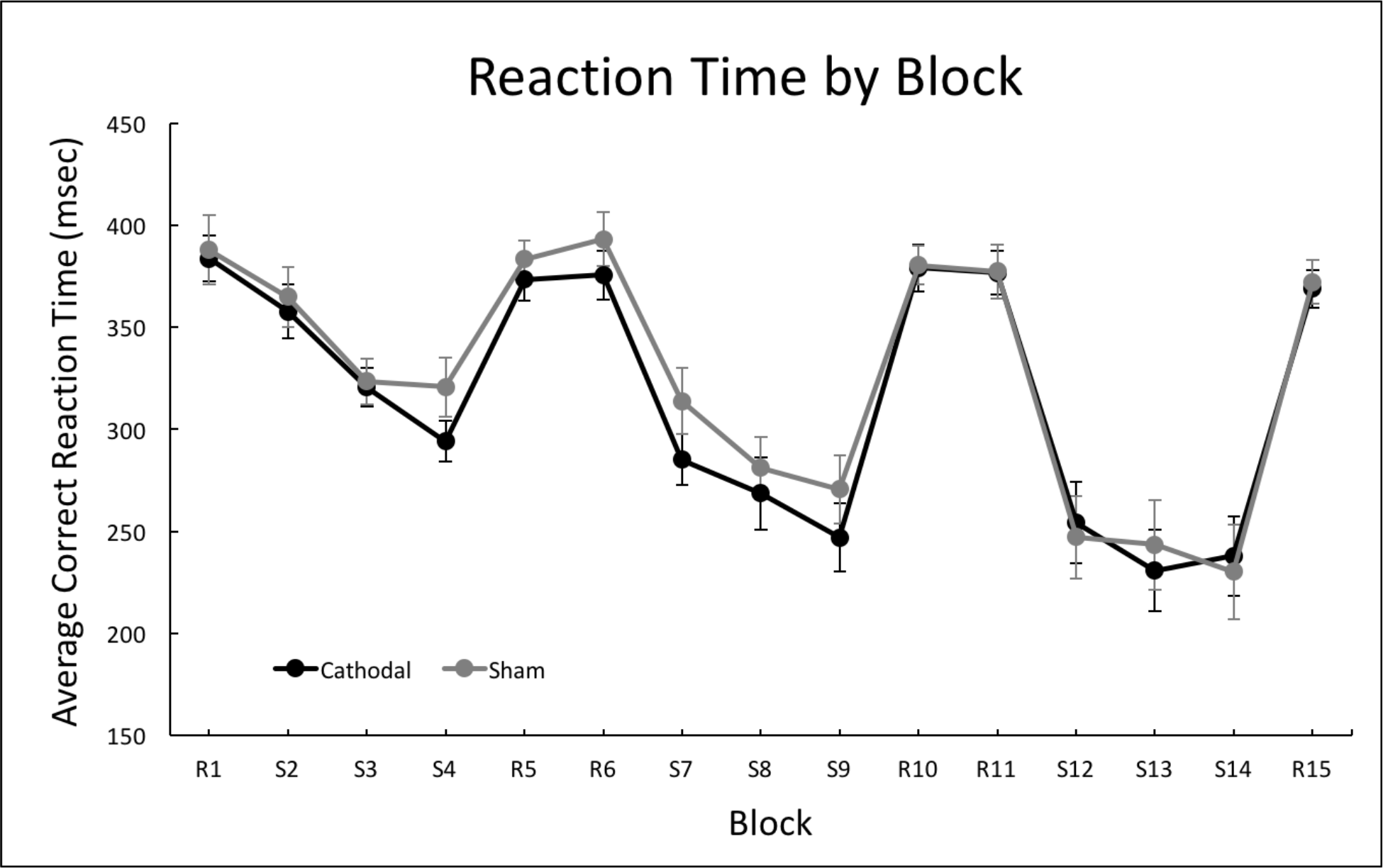
Group mean of correct reaction time, excluding missed trials, across all blocks in both cathodal and sham conditions for cerebellar stimulation. Error bars indicate standard error (SE).

#### Accuracy

A 3 × 2 repeated measures ANOVA (learning phase by stimulation condition), on sequence blocks only, revealed a significant learning phase by stimulation condition interaction, *F* (2, 40) = 3.35, *p* < .05, η^2^_p_ = .143. However, there was no significant main effect of learning phase, *F* (2, 40) = 0.919, *p* = 0.41, η^2^_p_ = .044, or stimulation condition, *F* (1, 20) = 1.58, *p* = 0.22, η^2^_p_ = .073. Follow-up paired samples *t*-tests to further unpack this interaction indicated that there is a marginally significant difference between cathodal and sham stimulation during the late phase of learning, *t* (20) = 2.07, *p* = .051. Accuracy during the early (*t* (20) = .119, *p* = .901) and middle (*t* (20) = .522, *p* = .607) phases of learning did not differ with stimulation. While individuals showed similar levels of performance during the first two phases of the task, performance was better (though only marginally significant) in the late phase, indicating an interaction between stimulation and performance depending on the specific phase of learning (Figure 3).

**Figure 3.**
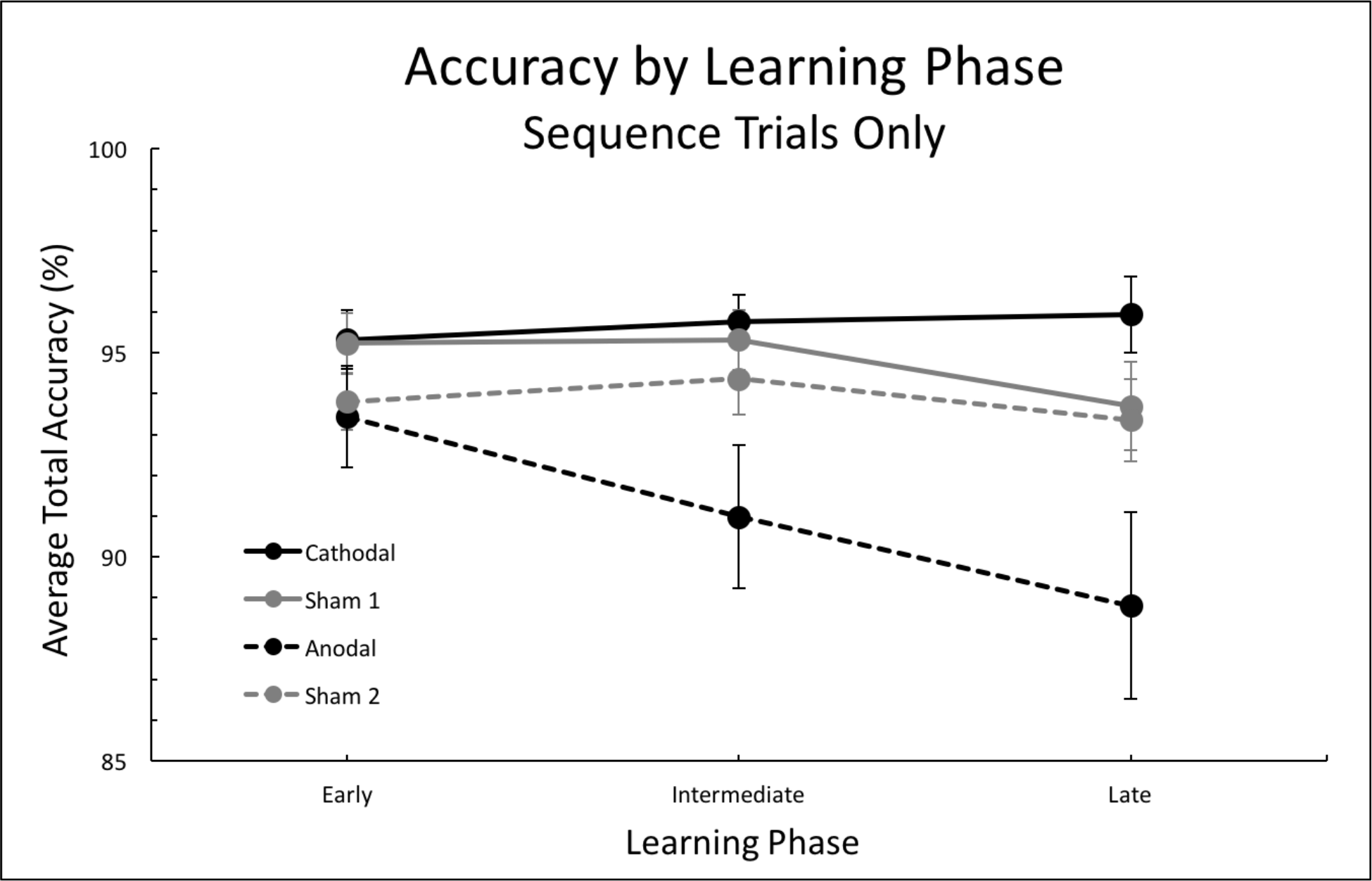
Group means of total accuracy, missed trials included, across early, intermediate, and late sequence learning phases in cathodal, sham-Experiment 1, sham-Experiment 2, and anodal conditions for cerebellar stimulation. Error bars indicate SE. Data from Experiment 1 is presented with solid lines, while that from Experiment 2 is presented with dashed lines.

## Experiment 2

### Materials and Methods

#### Participants

Thirty-five healthy young adults participated in the second experiment, consisting of two HD-tDCS sessions (anodal and sham) that were counterbalanced across subjects. Individuals in this experiment were completely separate from those in Experiment 1, and, as such, none of our subjects participated in both experiments resulting in two completely independent datasets. Three of those individuals did not return for the second session of the study, one individual did not undergo tDCS due to potential interactions with a medication, complete data was not collected from one individual due to technical difficulties, and one individual did not complete a full session due to discomfort from stimulation. Additionally, six individuals were excluded from final analyses for having task performance scores below three standard deviations from the group mean. The same *a priori* cut-off from Experiment 1 was applied here to uniformly exclude behavioral outliers. Those excluded performed the task with less than 46% accuracy. As such, our final sample included twenty-three subjects (15 female, mean age 18.6 years ± 0.9 S.D., range 18-21 years). None of the subjects had any history of medical or neurological disease and all were right-handed (mean score 93.9 ± 12.7 S.D.). Subjects were screened for the same exclusion criteria as in Experiment 1 (Nitsche et al., 2008), and none were taking medication that could potentially impact the central nervous system. Subjects were again recruited and scheduled through the Texas A&M University Psychology Subject Pool and received an equal amount of course credit as those in the first experiment. Participation was again limited to individuals that had not previously completed other tDCS studies. Prior to the initiation of study procedures, all subjects provided written informed consent, and the protocol was submitted to, and approved by, the Institutional Review Board at Texas A&M University.

#### Procedure

All procedures employed in the second experiment were identical to those described for Experiment 1, with the exception of the polarity of stimulation. Participants underwent two sessions, exactly one week apart, in which they received both anodal and sham tDCS in a within-subjects design. We again used a single-blind procedure wherein participants were not told which condition they were receiving, and the order of stimulation condition was counterbalanced. The electrode array and current intensities used for anodal cerebellar stimulation are presented in Table 2 and the associated modeled current flow is identical to that of Figure 1, but the flow of current is in opposite direction. This was achieved using a “spiral in” approach in this experiment, equivalent to anodal stimulation in a traditional two electrode pad system, again, determining the direction of current flow. Notably, the electrode placements and current intensities are identical to those in Experiment 1 in order to replicate the stimulation design with 2.0 mA of current applied to the lateral posterior cerebellum; however, the polarity of the current has been flipped. The same stimulation procedures from the first experiment were implemented in the second to avoid the subjects’ detection of stimulation condition and maintain consistency in our experimental design.

**Table 2.** Electrode array and current intensities for cerebellar anodal stimulation. *Location:* particular locations for electrode placement, determined by 10-20 system; *Current:* specific intensity of electrical current applied at a given location.

|  | <u>Location</u> | <u>Current</u> |
| --- | --- | --- |
| 1 | P2 | -0.1135 |
| 2 | PO10 | 0.1551 |
| 3 | O10 | 0.3618 |
| 4 | EX3 | -0.1956 |
| 5 | EX4 | 1.4832 |
| 6 | EX5 | -0.4066 |
| 7 | EX12 | -0.22 |
| 8 | EX14 | -0.7886 |
| C | NK1 | -0.2757 |

The motor sequence learning paradigm directly paralleled that from Experiment 1 (Kwak et al., 2012), and all parameters were identical. Participants were presented with a new sequence pattern during the second session of this experiment, and each sequence was comprised of 6 elements that did not contain any trills or sequential numbers. The same sequences used in the first experiment were employed here as well. As we had all new participants in Experiment 2, there was no concern with individuals already having learned the sequence.

#### Data Analysis

As in Experiment 1, performance was measured by average total accuracy (ACC) per sequence block and mean correct reaction time (RT) during both random and sequence blocks. Additionally, scores for each measure were calculated identically to those used in the first experiment. Repeated-measures ANOVAs and paired sample *t-*tests were performed to investigate performance differences between anodal and sham stimulation.

### Results

#### Reaction Time

A 2 × 2 repeated measures ANOVA (block type by stimulation condition), collapsing across all random and sequence blocks, revealed a significant main effect of block type on RT, *F* (1, 22) = 77.78, *p* < 0.001, η^2^_p_ = .78, wherein RT was significantly slower during the random blocks as compared to the sequence blocks. However, there was no significant main effect of stimulation condition (anodal vs. sham), *F* (1, 22) = 0.27, *p* = 0.61, η^2^_p_ = .012, or an interaction between block type and stimulation condition, *F* (1, 22) = 0.104, *p* = 0.74, η^2^_p_ = .005, consistent with Experiment 1. This indicates that participants perform the task more quickly when presented with sequence trials in comparison to random, as would be expected when movements are repeated in a sequential manner and said sequence is learned (Figure 4). Further, follow-up *t*-tests reveal a significant difference in reaction time between the first and last sequence blocks for this experimental group (*p* < .001). However, though reaction time improves over the course of the sequence blocks, accuracy does not improve, presumably due to the effects of stimulation on task performance.

**Figure 4.**
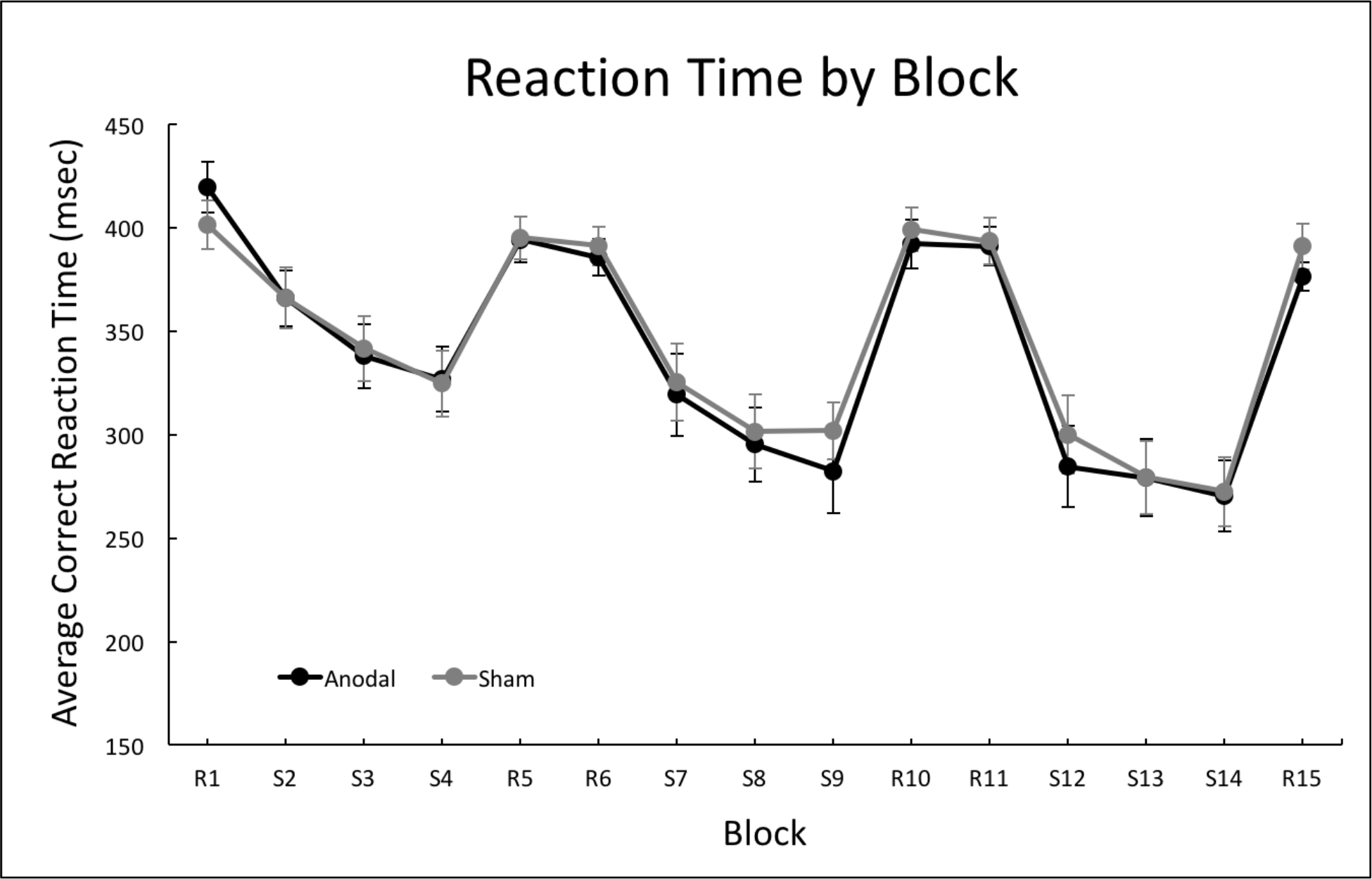
Group mean of correct reaction time, excluding missed trials, across all blocks in both anodal and sham conditions for cerebellar stimulation. Error bars indicate SE.

#### Accuracy

A 3 × 2 repeated measures ANOVA (learning phase by stimulation condition) revealed a significant learning phase by stimulation condition interaction (Figure 3), *F* (2, 44) = 3.43, *p* < .05, η^2^_p_ = .135, and a significant main effect of learning phase, *F* (2, 44) = 3.34, *p* < .05, η^2^_p_ = .132. The main effect of stimulation condition was approaching the standard cut-off for statistical significance, *F* (1, 22) = 3.65, *p* = .069, η^2^_p_ = .142. Follow-up paired samples *t*-tests to investigate this interaction revealed that there was no significant accuracy difference between sham and anodal stimulation during early learning (*t* (22) = − .353, p = .73), though there was a significant difference during the intermediate phase (*t* (22) = − 2.069, p = .05), and a difference that was approaching significance during the late phase of learning (*t* (22) = − 1.980, p = .06). In both the intermediate and late phases, performance following anodal stimulation was worse than that following sham stimulation; however, performance did not differ during early learning, driving the interaction. Thus, though participants are responding more quickly, they are doing so with poorer accuracy over time. Performance is getting worse as assessed by accuracy.

## Comparing Cathodal and Anodal Stimulation

In order to follow up on the results from Experiments 1 and 2 and get a better idea of the effect of stimulation type on explicit motor sequence learning, we conducted additional between-subjects analyses on the two independent groups. These statistical procedures were carried out as *a priori* analyses aimed at accomplishing a comprehensive perspective on the nature of the cerebellum’s role in motor learning by comparing the stimulation conditions. In all cases, we used 3 × 2 (learning phase by stimulation group) mixed model ANOVAs. Because significant stimulation effects were limited to the accuracy measure, we focused our analyses on accuracy. First, we investigated group differences on the sham condition only, to determine whether or not there were any baseline learning differences between the two experimental groups. Critically, there was no significant main effect of group, *F* (1, 42) = .796, *p* = .377, η^2^_p_ = .019. Further, there was no significant interaction between learning phase and group, *F* (2, 84) = .441, *p* = .645, η^2^_p_ = .010, though the main effect of phase was approaching significance, *F* (2, 84) = 2.73, *p* = .071, η^2^_p_ = .061. Therefore, in the absence of stimulation, participants exhibited similar degrees of performance on the sequence trials across both sham groups.

Next, we directly compared performance between the two groups during active stimulation (cathodal compared to anodal). This revealed a significant learning phase by group interaction, *F* (2, 84) = 4.35, *p* < .05, η^2^_p_ = .094, as well as a significant main effect of group, *F* (1, 42) = 6.79, *p* < .05, η^2^_p_ = .139. The main effect of learning phase was approaching significance, *F* (2, 84) = 2.53, *p* = .085, η^2^_p_ = .057. As seen in Figure 3, participants perform significantly better on the sequence learning paradigm after cathodal cerebellar stimulation compared to anodal. Thus, stimulation has polarity-specific effects on our explicit motor sequence learning paradigm.

Further, we calculated difference scores relative to sham (i.e., ACC cathodal group – ACC sham Experiment 1 group and ACC anodal group – ACC sham Experiment 2 group) for both experiments, so as to compare across groups relative to baseline. This exploratory, *post hoc* analysis was implemented to provide information about the magnitude of learning achieved within each group after active stimulation when different conditions were compared both directly and also in consideration of baseline performance after sham stimulation. Importantly, as described above, accuracy performance between the groups in the absence of stimulation did not differ, justifying our use of the sham scores as controls for this specific measure. First, there was a significant learning phase by stimulation group interaction, *F* (2, 84) = 5.42, *p* < .01, η^2^_p_ = .114. There was also a significant main effect of group, *F* (1, 42) = 4.88, *p* < .05, η^2^_p_ = .104, though the main effect of learning phase was not significant, *F* (2, 84) = 1.04, *p* = .36, η^2^_p_ = .024. Following up on these analyses, independent samples *t*-tests revealed that the interaction is driven by a significant difference in performance relative to sham during late learning specifically, *t* (42) = 2.59, *p* = .01, and a significant difference during the middle phase, *t* (42) = 2.02, *p* = .05, though there were no differences during early learning, *t* (42) = .35, *p* = .728. This provides further support for polarity-specific effects of cerebellar stimulation on explicit motor sequence learning, even when normalized relative to baseline (sham stimulation). Cathodal stimulation is resulting in improvement whereas anodal is making performance worse over time, specifically when scores from each active condition are normalized to sham and subsequently compared.

## Discussion

Applying HD-tDCS to the lateral posterior cerebellum prior to the learning of a new motor sequence may have helped facilitate initial skill acquisition when using cathodal stimulation (relative to sham), whereas anodal stimulation negatively impacted performance (relative to sham). Across the learning phases, accuracy increased with cathodal stimulation (though these increases were quite small) but decreased with anodal when comparing between experiments. Critically, performance across the two experiments and stimulation types significantly differed, suggesting a polarity-specific effect of stimulation on accuracy during explicit motor sequence learning. However, we did not see any effects on RT other than differences in speed between block types, wherein responses were significantly faster during sequence blocks relative to random blocks, across both experiments. These findings and their implications are discussed, in turn, below.

With respect to performance accuracy, we found that cathodal cerebellar stimulation provides a benefit relative to sham. Specifically, this was driven by improved performance towards the later stages of our paradigm, though only marginally significant. Our results suggest that cathodal stimulation is more favorable than no stimulation at all, but the modest nature of these within-subject findings from Experiment 1 do not conclusively indicate that we have directly facilitated learning. Rather, we may have prevented any potential fatigue or attentional declines that may occur during the end of the task. Nonetheless, cathodal cerebellar stimulation positively impacted sequence learning. Conversely, anodal stimulation had a negative influence on skill acquisition. While performance was nearly identical to that of the sham condition during the first few blocks of learning (here, the early phase), performance after anodal stimulation was significantly worse later in the task. These effects were considerably more pronounced in the anodal group compared to the cathodal, but the between-subject analyses further supported the idea that a differential influence on learning is observed across the two conditions. As such, these results indicate polarity-specific effects of tDCS on explicit motor sequence learning.

As this paradigm is only performed once and without extensive practice, these effects take place during the overall earlier stages of explicit sequence learning when cognitive resources, such as working memory, are put to use. Stimulation was specifically applied to a cognitive region of the cerebellum, namely the lateral posterior cerebellum (roughly including Crus I and II), which has been implicated in working memory processes (Schmahmann and Sherman, 1998; Stoodley, 2012; Bernard and Seidler, 2013, 2014; Guell et al., 2018). This may, therefore, explain the early influence on motor skill acquisition as the motor pattern is being actively rehearsed and held at the forefront of the mind in light of explicit information. Further, it is possible that our stimulation montage exerted some downstream effects on the prefrontal cortex as the cerebellum maintains closed-loop connections with this cognitive area (Krienen and Buckner, 2009; O’Reilly et al., 2010; Salmi et al., 2010; Bernard et al., 2012, 2014). We speculate that the diverging impacts on accuracy that emerged over the course of the task after stimulation is due to these networks and functions. That is, cathodal stimulation of the lateral posterior cerebellum may have “primed” or activated network circuits involved in working memory, allowing them to be more effectively utilized when cognitive demands are higher in these initial skill acquisition phases of motor sequence learning. It is possible that our approach may have influenced the cognitive processes at play more directly (Desmond et al., 2005; Oliveri et al., 2007; Ferrucci and Priori, 2014), but there is also evidence to suggest that cerebellar stimulation induces functional network changes that may underlie the observed effects as well (Halko et al., 2014). Cathodal stimulation may have facilitated working memory resources and the associated cerebellar circuits that communicate with the cerebral cortex. In contrast, anodal stimulation may have inhibited these cerebellar cortical regions and circuits, making it more difficult for participants to engage cognitive resources, presumably verbal working memory (cf. Bo and Seidler, 2009), during this overall earlier learning period.

Modulating, and specifically improving, motor learning is an area of great interest, given the potential implications that success in this regard would have for rehabilitation after injury or infarct. For a broad review on the effects of tDCS on motor learning, we refer readers to Buch et al. (2017). Pertinent to our work here, there have been several studies targeting the cerebellum specifically (e.g., Galea et al., 2011; Block and Celnik, 2013; Foerster et al., 2013; Shah et al., 2013; Hardwick and Celnik, 2014; Cantarero et al., 2015), though these investigations have primarily focused on adaptation tasks. As such, our investigation here using explicit sequence learning presents a novel extension of this literature as a whole. Across multiple investigations of adaptation learning, results suggest that anodal tDCS can facilitate motor learning (Galea et al., 2011; Block and Celnik, 2013; Hardwick and Celnik, 2014), though cathodal stimulation has also been shown to facilitate learning in an ankle adaptation task (Shah et al., 2013). More recently, work from Cantarero et al. (2015) included both cathodal and anodal conditions, during which individuals performed a task involving the production of isometric force with a design that includes sequencing components. Notably, they found polarity specific effects, wherein anodal stimulation improved performance, while cathodal stimulation had no effect on learning.

In the present study, we found polarity specific effects of stimulation. However, the directionality of these findings differed from the larger literature using tDCS to modulate motor learning. Our results suggest that cathodal stimulation may have had a small facilitatory effect, whereas anodal stimulation had a negative effect on learning, as measured by accuracy. While at first this stands in contrast to the given literature, there are several key considerations. First, the adaptation work has largely relied upon online tDCS stimulation (Galea et al., 2011; Block and Celnik, 2013; Cantarero et al., 2015). That is, stimulation was applied during performance of the learning paradigm. Cantarero et al. (2015) suggest that this online stimulation may aid in the processing of motor errors in the cerebellar cortex. Second, our investigation took advantage of an HD-tDCS system, whereas work to date has primarily relied upon two electrode stimulation pads (Ehsani et al., 2016; Wessel et al., 2016; Shimizu et al., 2017). In addition, these studies have employed different types of stimuli and made use of task designs distinct from our own, as the ultimate intention of these investigations was to examine motor retention rather than skill acquisition as is our objective in the current study. Thus, differences in the underlying methodology and experimental aims may contribute to these discrepancies as well. However, Shah et al. (2013) did see comparable facilitation with both anodal and cathodal stimulation, and a recent meta-analysis of the effects of cerebellar tDCS suggests that while stimulation impacts behavior (both motor and cognitive) across multiple studies, there is little support to suggest that these effects are polarity specific (Oldrati and Schutter, 2018). Finally, the underlying physiology of the cerebellum is also of note.

In an investigation of the impact of cerebellar tDCS on cognition, Pope et al. (2012) demonstrated that cathodal stimulation resulted in behavioral improvements, while anodal stimulation had no impact on behavior. This is broadly consistent with our findings here, and our goal was to ultimately modulate cognitive networks and systems. Notably, Pope et al. (2012) suggest that this seemingly counterintuitive result is due to the inhibitory action of cerebellar Purkinje cells on the deep cerebellar nuclei, which are the primary output regions of this structure. Because Purkinje cells have an inhibitory effect on the deep cerebellar nuclei, presumably the dentate nucleus in this particular investigation, cathodal stimulation may decrease the action of the Purkinje cells and release this inhibition (Pope and Miall, 2012). Reduced inhibition may in turn result in facilitation of cerebello-cortical networks.

This is also consistent with the idea of cerebellar-brain inhibition. With transcranial magnetic stimulation, stimulation to the cerebellum results in inhibition over the primary motor cortex (Popa et al., 2010), and when tDCS is used, there are comparable polarity-specific effects (Galea et al., 2009). Cathodal tDCS to the cerebellum decreases this inhibition, while anodal stimulation increases it. While this work has focused on the primary motor cortex, we speculate that the comparable circuitry with the prefrontal cortex acts in a similar manner. Thus, in the present study, we suggest that stimulation to the lateral posterior cerebellum results in either an increase in or release from inhibition on prefrontal cortical regions and networks depending on stimulation polarity, paralleling what was suggested by Pope et al. (2012). This would then result in either negative impacts or relative improvement on accuracy during explicit motor sequence learning. Targeting these cerebello-thalamo-frontal circuits may have inhibited or facilitated the use of working memory resources, which in turn influenced accuracy during the learning of a motor pattern. Future work incorporating functional neuroimaging methodologies, however, is needed to further investigate this idea.

More broadly, our findings further underscore the importance of the lateral posterior cerebellum in non-motor behavior and suggest that the cerebellum may also contribute to the cognitive aspects of initial skill acquisition. In addition to contributing to the control of movements necessary to complete the task, we suggest that cerebellar networks associated with cognitive processing are also used during learning. Prior work has linked motor sequence learning with working memory capabilities (Bo and Seidler, 2009), suggesting that those with higher working memory capacity are better able to chunk sequence elements. Notably, the cortical networks that typically support sequence learning include the basal ganglia, prefrontal cortex, parietal cortex, and of course, primary motor cortex (Hikosaka et al., 1999; Doyon et al., 2002; Seidler, 2006; Galea et al., 2011). While the cerebellum is also activated during sequence learning, it is less prevalent, particularly as compared to activation seen during adaptation learning paradigms (Baizer et al., 1999; Imamizu et al., 2000, 2003; Anguera et al., 2010). However, as previously noted, the region we targeted with stimulation is associated with working memory processes as well as language (Stoodley and Schmahmann, 2009; Stoodley, 2012). Stimulation may have therefore facilitated the cortical networks necessary for efficient chunk formation (though we did not directly measure chunking here) and allowed for the use of verbal working memory rehearsal strategies during initial skill acquisition.

While our findings provide interesting new insights into the role of the cerebellum in explicit sequence learning, there are several limitations to consider. First, we stimulated the cerebellum and suggest that we may have facilitated activation in the broader cerebello-prefrontal network associated with working memory (Krienen and Buckner, 2009; Stoodley and Schmahmann, 2009; Bernard et al., 2012; Stoodley, 2012); however, this stimulation may have spread to other regions of the cerebellum. Though HD-tDCS allows for improved targeting, the stimulation does reach other areas that could have an effect on performance (Figure 1). And further, the cerebellum is also known to play a key role in timing (Ivry and Keele, 1989; Ivry et al., 2002; Spencer and Ivry, 2013). While typically the more medial regions of the cerebellum have been implicated in this function (Ivry et al., 1988), again, the spread of the stimulation effect may have helped facilitate timing. Improved event and movement timing may, in turn, result in improved accuracy in initial skill learning. To further test our suggestion that improved working memory impacts explicit sequence learning, stimulation of the prefrontal cortex in future work is warranted as well. Second, we suggest that this working memory facilitation may also aid in the chunking of motor elements (Bo and Seidler, 2009). However, we did not explicitly measure chunking as the employed sequences were relatively short and our inter-stimulus interval was fairly small and fixed, so this, therefore, remains speculative. Relatedly, individuals may have potentially approached the task with different strategies in each experiment, but the lack of performance differences across the two sham groups and the counterbalancing implemented in our design likely reduces this possibility. Additionally, the long-term benefits of stimulation are not clear. Our sequence learning paradigm was fairly brief, and we did not measure retention over time. Thus, it would be beneficial in future work to investigate more long-term differences in performance after cathodal cerebellar stimulation compared to anodal and relative to sham.

Due to the nature of working with participant subject pools, subject attrition can be an issue. We therefore chose to conduct two separate experiments with independent samples and then compare across stimulation conditions as a fully within-subjects design is difficult to accomplish with the recruitment opportunities available, though such a design would be optimal. This constraint also led us to exclude a considerable number of subjects that did not complete a second session so that we could conduct our analyses on a full, balanced dataset. Moreover, our subjects skew female, and this is an artifact of the sample from which our participants were selected. Though we are not able to reliably test any possible gender effects due to power constraints, this may have impacted our findings. Finally, while we generally see that anodal cerebellar tDCS negatively impacts performance, the extent to which this is the case is widely variable, as demonstrated by the large error bars seen in Figure 3. This may, in part, be due to a considerable degree of variability in responsiveness to stimulation as the impact on performance can fluctuate with respect to individual differences (Chew et al., 2015; Li et al., 2015).

## Conclusions

Together, these findings provide key new insights into the role of the cerebellum in explicit sequence learning. Our results suggest that stimulating the lateral posterior cerebellum with a cathodal approach benefits initial skill acquisition, relative to both sham and anodal stimulation. In addition, our results suggest that, compared to both sham and cathodal stimulation, applying anodal tDCS to this particular region of the cerebellum negatively impacts explicit motor sequence learning. Therefore, these findings advance our understanding of the role of the cerebellum in explicit sequence learning as we suggest that we may have influenced cognitive processes in verbal working memory, often relied upon during initial learning. Further, exploring techniques that may improve motor skill acquisition is valuable for the development of new therapeutic targets for neurological disorders (e.g., Parkinson’s disease), as well as enhancing restorative therapy after stroke or other injuries impacting motor function and reducing normative motor declines with advanced age.

